# AtDjB3 regulates Hsc70-1-mediated expression of the heat shock genes and thermotolerance in Arabidopsis

**DOI:** 10.1101/2025.05.20.654906

**Authors:** Gouri Satheesh, Ishani Sengupta, Chandan Sahi

## Abstract

Heat stress disrupts protein homeostasis, triggering the heat shock response (HSR) to maintain cellular proteostasis. A key aspect of this response is the release of heat shock factors (Hsfs) from Hsp70-mediated attenuation under heat-shock conditions, making Hsp70 a central regulator of HSR. However, the role of Hsp70 co-chaperones in this process remains largely unexplored in plants. Our study identifies AtDjB3, a heat-inducible class II J-domain protein (JDP), as a critical modulator of HSR. We present microscopy and cell fractionation-based evidence to demonstrate that the loss of AtDjB3 impairs Hsc70-1 recruitment to heat-induced protein aggregates, thereby maintaining HsfA1d bound to Hsc70-1 in the cytoplasm. This inhibited the expression of many heat-inducible genes, including *HSP70s*, *HSP90*, *HSP18.2*, *FES1A*, *HSFA2*, and *HSFA7A*. Consistent with this, *AtDJB3* mutants displayed compromised thermotolerance, as evidenced by their inability to survive a prolonged heat stress of 37°C. Conversely, overexpression of *AtDJB3* conferred enhanced thermotolerance, further supporting its positive regulatory role in the HSR. We propose that AtDjB3 not only contributes to the solubilization of heat-induced protein aggregates but also promotes Hsf activation by diverting Hsc70-1 to these aggregates, thereby releasing Hsfs to drive the transcriptional heat shock response.

## Introduction

Hsp70-JDP (J-domain protein) machines are crucial for mitigating stress-induced damage across all organisms (Alagar Boopathy et al., 2022; Kityk et al., 2018). They operate by facilitating protein folding, preventing aggregation, and aiding in the refolding or degradation of misfolded proteins (Kampinga and Craig, 2010; Kohler and Andréasson, 2020). Additionally, Hsp70 has a key role in activating heat shock response (HSR), a transient, highly coordinated, and conserved transcriptional response that enables cells to rapidly activate the expression of several stress-associated genes (Hahn et al., 2011; Ohama et al., 2017). Activation of HSR is fundamental to an organism’s ability to endure and adapt to elevated temperatures (Bakery et al., 2024; Mittler et al., 2012).

HSR is controlled by the rapid activation of heat shock transcription factors (Hsfs). Arabidopsis encodes 21 Hsfs classified into three distinct groups: A, B, and C, based on the characteristics of their flexible linkers and oligomerization domains (Nover et al., 2001; Scharf et al., 2012). Among them, HsfA1s and HsfA2 function as key positive regulators of HSR (Charng et al., 2007; Liu et al., 2011; Schramm et al., 2006; Von Koskull-Döring et al., 2007; Yoshida et al., 2011). Consistent with their central role in regulating HSR and plants response to high temperature, a quadruple mutant of HsfA1s (*hsfa1a*, *hsfa1b*, *hsfa1d*, *hsfa1e*) in Arabidopsis displays severely diminished expression of most heat-induced genes and compromised thermotolerance (Liu et al., 2011). Studies in other models, including Chlamydomonas, rice, and tomato have shown that HsfA1s are constitutively expressed under non-stress conditions and rapidly upregulated upon heat stress to activate heat-responsive genes and promote thermotolerance (Liu et al., 2011; Mishra et al., 2002; Yoshida et al., 2011). Among others, HsfA1d plays a particularly significant role in regulating heat-induced gene expression (Kotak et al., 2007; Nishizawa-Yokoi et al., 2011; Yoshida et al., 2011). Intriguingly, some Hsfs, such as HsfB1 and HsfB2b, function as transcriptional repressors (Ikeda et al., 2011), making genetic and functional studies on Hsfs immensely complicated. While HsfA1s are widely recognized as master regulators of HSR, they also contribute to broader environmental stress responses, plant growth, development, as well as reproductive fitness (Guo et al., 2016; Haider et al., 2022; Majee et al., 2023).

Hsp70s play a dual role in heat stress adaptation. In addition to mitigating heat-induced proteostasis defects, Hsp70 also modulates HSR through direct physical interactions with Hsfs (Hahn et al., 2011; Tiwari et al., 2020; Usman et al., 2017). Hsp70 negatively regulates Hsf activity across species through a feedback mechanism to control its expression and that of other heat-responsive genes (Ciccarelli and Andréasson, 2024; Kim and Schöffl, 2002; Masser et al., 2019). In Arabidopsis, *HSC70-1* mutants exhibit increased transcript levels of several heat-responsive genes during and after heat stress, enhancing basal thermotolerance. Conversely, *HSC70-1* overexpression results in a thermosensitive phenotype (Tiwari et al., 2020).

Although Hsp70 is a key molecular chaperone, its function is largely dependent on co-chaperones known as J-domain proteins (JDPs), which determine their specificity and function (Kampinga et al., 2019; Usman et al., 2017). Among JDPs, Class II/B-type JDPs have been implicated in HSR in various model organisms. These proteins contain an N-terminal J-domain, a glycine-rich (G/F) region, and two C-terminal β-barrel domains (CTD I and CTD II), which house the substrate-binding region, followed by a dimerization domain (DD) (Lee et al., 2002). Class II JDPs, such as DroJ1 in Drosophila and DnaJB1 in humans, regulate HSR by directly interacting with Hsf, promoting its binding to Hsp70, and inducing conformational changes in Hsf1 through ATP-dependent Hsp70 activity (Kmiecik et al., 2020; Marchler and Wu, 2001). In budding yeast, the Class II JDP Sis1 mediates substrate delivery to Hsp70 during heat stress, leading to Hsf1 dissociation from Hsp70 and subsequent activation of HSR (Feder et al., 2020). These mechanistic variations underscore the functional versatility of Class II JDPs while reinforcing their central role in protein homeostasis and stress adaptation.

In *Arabidopsis thaliana*, eight cytosolic Class II JDPs (AtDjB1, AtDjB2, AtDjB3, AtDjB4, AtDjB5, AtDjB6, AtDjB10, and AtDjB17) have been identified (Verma et al., 2019, 2017). AtDjB3, in particular, has been recognized for its involvement in acquired thermotolerance and the resolubilization of protein aggregates formed under heat stress (Tak et al., 2023). However, their role in the regulation of heat-induced gene expression is not known. By integrating phenotypic, molecular, and biochemical approaches, this study identifies a novel regulatory mechanism in which AtDjB3, in collaboration with Hsc70-1, ensures efficient HSR. We demonstrate that AtDjB3 is required for the effective recruitment of Hsc70-1 to heat-induced protein aggregates, thereby promoting the nuclear translocation of HsfA1d and activating HSR. These findings reveal a critical interplay between chaperones and transcription factors in plant stress responses, offering new insights into the molecular mechanisms underlying thermotolerance.

## Results

### AtDjB3 protein accumulates under high temperatures and associates with heat-induced protein aggregates

AtDjB3 transcription is upregulated in response to heat stress (Tak et al., 2023). To evaluate whether AtDjB3 protein abundance mirrors this transcriptional upregulation, a stable GFP-tagged transgenic line of AtDjB3 expressed from its own promoter (*pAtDJB3::AtDJB3- GFP*) was generated. Ten-day-old seedlings were exposed to temperatures ranging from 22°C to 42°C for 2h. Immunoblotting showed significant induction of AtDjB3-GFP fusion protein, with noticeable accumulation at 35°C, with expression peaking at 37°C. Interestingly, AtDjB3 levels declined at 40°C, indicating a narrow range of heat-inducibility of AtDjB3 protein **(Fig. 1A)**. When plants were exposed to 37°C for 24h, AtDjB3 protein became detectable within 30 min, reached maximal levels by 4h, and gradually declined thereafter **(Fig. 1B)**. These results demonstrate that AtDjB3 protein expression is rapidly and consistently expressed during heat stress, suggesting a possible role in orchestrating heat stress response in Arabidopsis.

**Figure 1.**
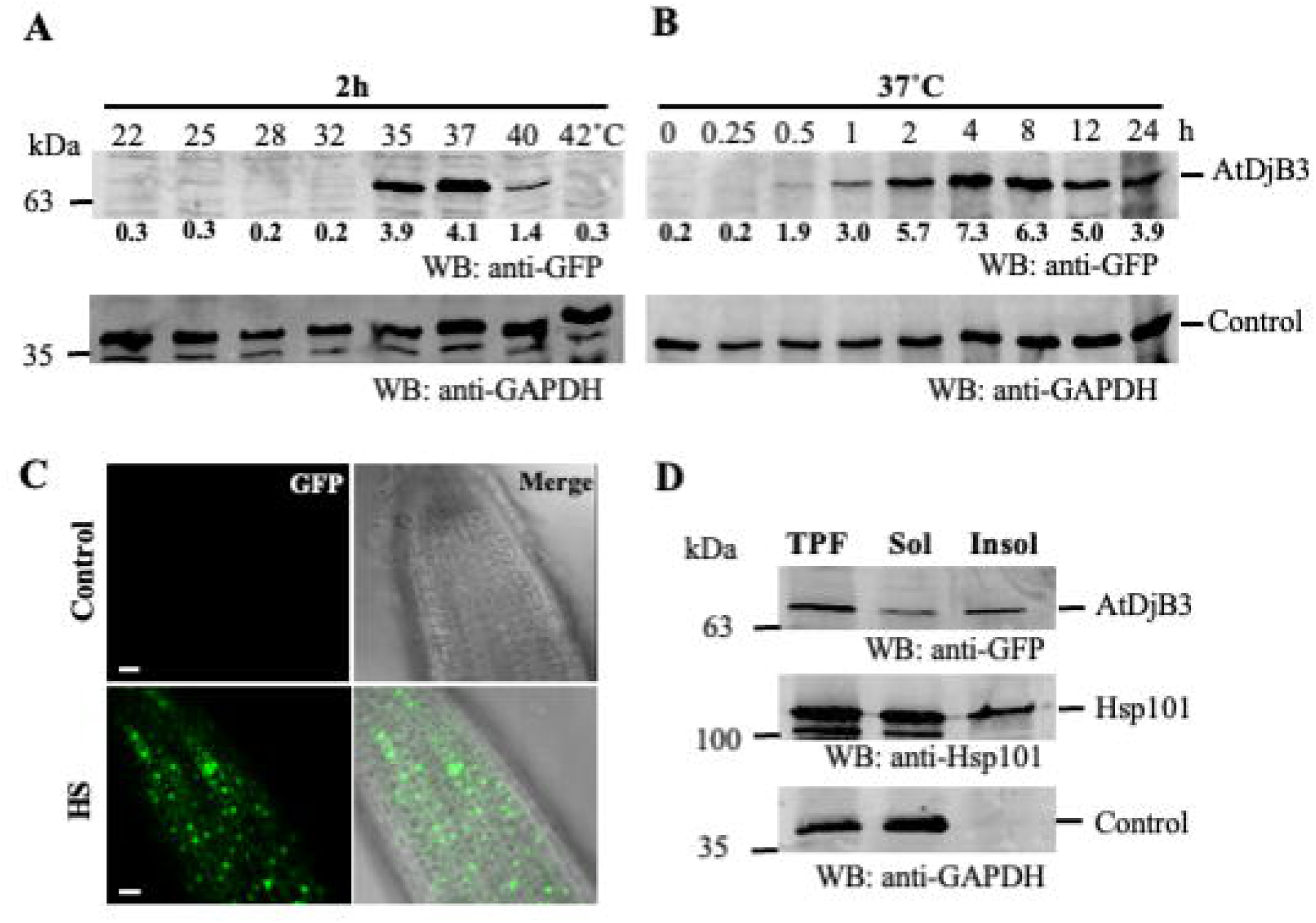
AtDjB3 protein is heat-inducible and associates with heat-induced protein aggregates. (A, B) Immunoblot analysis of total protein extracts from I O-day-old AtDjB3-GFP seedlings subjected to cither different temperatures for 2h or to 37°C for various durations. Blots were probed with anti-GFP; GAPDH served as a loading control. Signal intensities were normalized to GAPDH. (C) Confocal imaging of roots from I O-day-old AtDjB3-GFP seedlings before and after 15-min heat stress at 37°C. *n* = 3. Scale bars. 10 pm. (D) Immunoblot of total, soluble, and insoluble protein fractions from seedlings subjected to 2h heat stress at 37°C. Blots were probed with anti-GFP; HsplOl and GAPDH were used as aggregate and loading controls, respectively. All immunoblotting experiments were performed in at least three independent biological replicates.

Previous studies have shown that AtDjB3 associates with heat-induced protein aggregates and when expressed from a strong constitutive promoter, colocalizes with protein aggregate centers (PACs) in Arabidopsis protoplasts (Tak et al., 2023). So, we examined the localization of AtDjB3-GFP when expressed from the native AtDjB3 promoter. When subjected to heat stress, AtDjB3-GFP showed punctate distribution within root cells of 10- day old Arabidopsis, a pattern reminiscent of PACs **(Fig. 1C, bottom)**. As expected, no GFP signal was detected under control conditions, confirming the tight heat-inducibility of AtDjB3 **(Fig. 1C, top)**. Fractionation analysis also confirmed that AtDjB3-GFP partitions into the insoluble fraction, along with Hsp101, a known marker for protein aggregates in Arabidopsis during heat stress **(Fig. 1D)** (McLoughlin et al., 2016). As expected, GAPDH was exclusively detected in the soluble fractions (McLoughlin et al., 2016; Tak et al., 2023). These results confirm that AtDjB3 is a heat-responsive co-chaperone that dynamically redistributes to protein aggregates, reinforcing its potential role during heat stress.

### AtDjB3 modulates thermotolerance in Arabidopsis

Given the rapid induction of AtDjB3 during heat stress, we explored its functional significance using T-DNA insertion mutants (*b3-1* and *b3-2*) and constitutive overexpression lines, all validated for homozygosity and AtDjB3 transcript levels **(Supplementary** Fig. 1). After a short-term heat stress (37°C, 2h), WT, *AtDJB3* T-DNA insertion mutants, and overexpression lines exhibited comparable, as indicated by the emergence of true leaves during the post-stress period **(Supplementary** Fig. 2). However, under prolonged heat stress of 37°C for 48h, *b3-1* and *b3-2* mutants displayed significant heat stress sensitivity, with nearly 98% of seedlings failing to recover compared to approximately 50% lethality in WT and complemented lines **(Fig. 2A, B)**. This was further supported by reduced fresh weight and chlorophyll content in the mutants compared to WT **(Fig. 2C, D)**. Growth of all the lines analyzed was comparable to Col-0 before the heat treatment **(Supplementary** Fig. 3). In contrast, AtDjB3 overexpression lines exhibited improved survival of about 85% following the prolonged stress, suggesting that enhanced AtDjB3 expression confers greater thermotolerance, further highlighting its critical role in high temperature stress tolerance.

**Figure 2.**
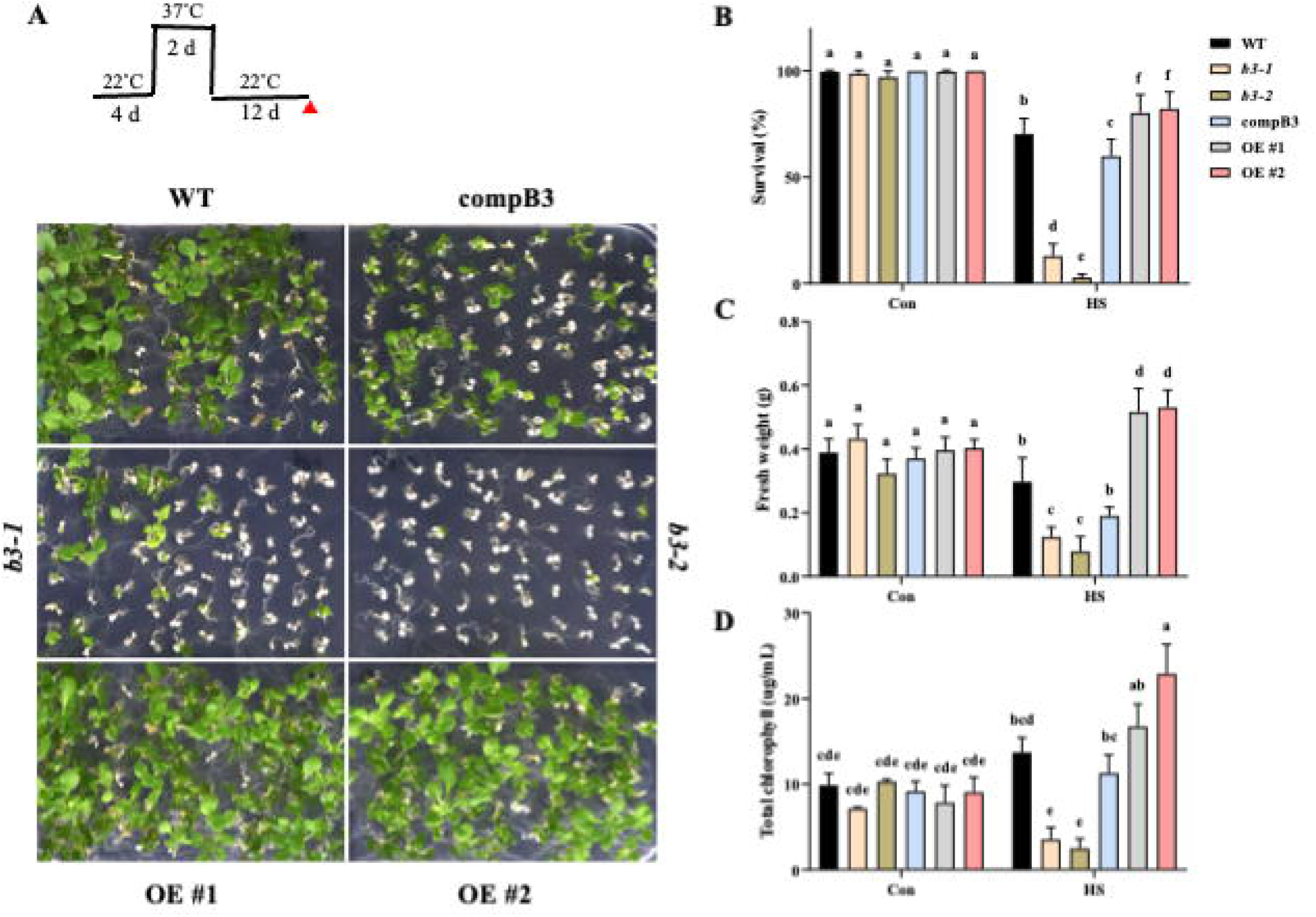
*AtDJB3* is critical for basal thermotolerance in Arabidopsis seedlings. (?\) 4-day-old seedlings of WT, *AtDJB3* mutants *(b3-l* and *b3-2),* complemented, and overexpression lines (OE #1 and #2) were subjected to 37°C for 2 days and allowed to recover under control conditions. Photographs were taken on day 12 post­treatment. (B-D) Survival rate, fresh weight, and total chlorophyll content were quantified across genotypes. Significant differences between groups arc determined using two-way ANOVA followed by Tukey’s HSD test *( P < 0.05).* Different lowercase letters denote statistically distinct groups. Data represent mean ± SE from four biological replicates *(N* = 150 seedlings).

### AtDjB3 is required for transcriptional activation of heat-responsive genes

AtDjB3 plays a crucial role in solubilizing heat-induced protein aggregates and is thereby required for acquired thermotolerance in Arabidopsis (Tak et al., 2023). A newfound role of AtDjB3 in basal thermotolerance prompted us to investigate the underlying mechanisms behind this phenomenon. We hypothesized that, like its orthologs, Sis1 in budding yeast (Feder et al., 2020), DroJ1 in Drosophila (Marchler and Wu, 2001), and DnaJB1 in humans (Kmiecik et al., 2020), AtDjB3 may also be involved in regulating HSR. To test this, we examined the expression of several heat stress-responsive genes in *AtDJB3* mutants, complemented, and overexpression lines. The *b3-1* mutant, extensively characterized in our previous studies, and the highly overexpressing *AtDJB3* line (OE #2) were chosen for transcriptional analysis. A subset of heat-responsive genes, including heat shock proteins (Hsps) and non-heat shock proteins (non-Hsps), was analyzed for transcriptional changes in response to high temperatures. Unlike the wild-type plants, in the *b3-1* line, the transcripts of several Hsps such as *HSC70-1*, *HSP70-2*, *HSP70-4*, *HSP70-5*, *HSP90-1*, small Hsps (*HSP17.6* and *HSP18.2*), *AtDjA1* (JDP), and *HSA32* were not upregulated under HS conditions. Additionally, transcripts encoding *MBF1C* (*MULTIPROTEIN BRIDGING FACTOR 1C*), *ROF2* (a peptidyl-prolyl cis-trans isomerase), *HSFA2*, *HSFA7A*, *HOP3*, and *FES1A* (a cytosolic NEF) were significantly downregulated in *b3-1* compared to WT **(Fig. 3)**, suggesting that *AtDJB3* is required for the activation of a few selected heat-inducible genes. However, genes, like *ROF1*, *AtDJA2* (JDP), *APX2* (*ASCORBATE PEROXIDASE 2*), *DREB2A* (*DEHYDRATION-RESPONSIVE ELEMENT BINDING PROTEIN 2A*), and *HSP70-3,* maintained expression levels similar to WT under heat stress conditions. To further investigate the regulatory function of *AtDJB3* in heat-responsive transcription, we examined its overexpression lines. Several genes, such as *HSC70-1*, *HSP70-4*, *HSP70-5*, *HSP17.6*, *HSP18.2*, *AtDJA1*, *HSA32*, *FES1A*, *ROF2*, and *MBF1C*, exhibited significantly elevated expression relative to WT **(Fig. 3)**, while *HSP90-1* and *HSP70-2* remained unchanged. These results highlight *AtDJB3* as a critical regulator required for the transcriptional activation of a subset of heat-inducible genes, reinforcing its role in modulating the heat shock response in *Arabidopsis thaliana*.

**Figure 3.**
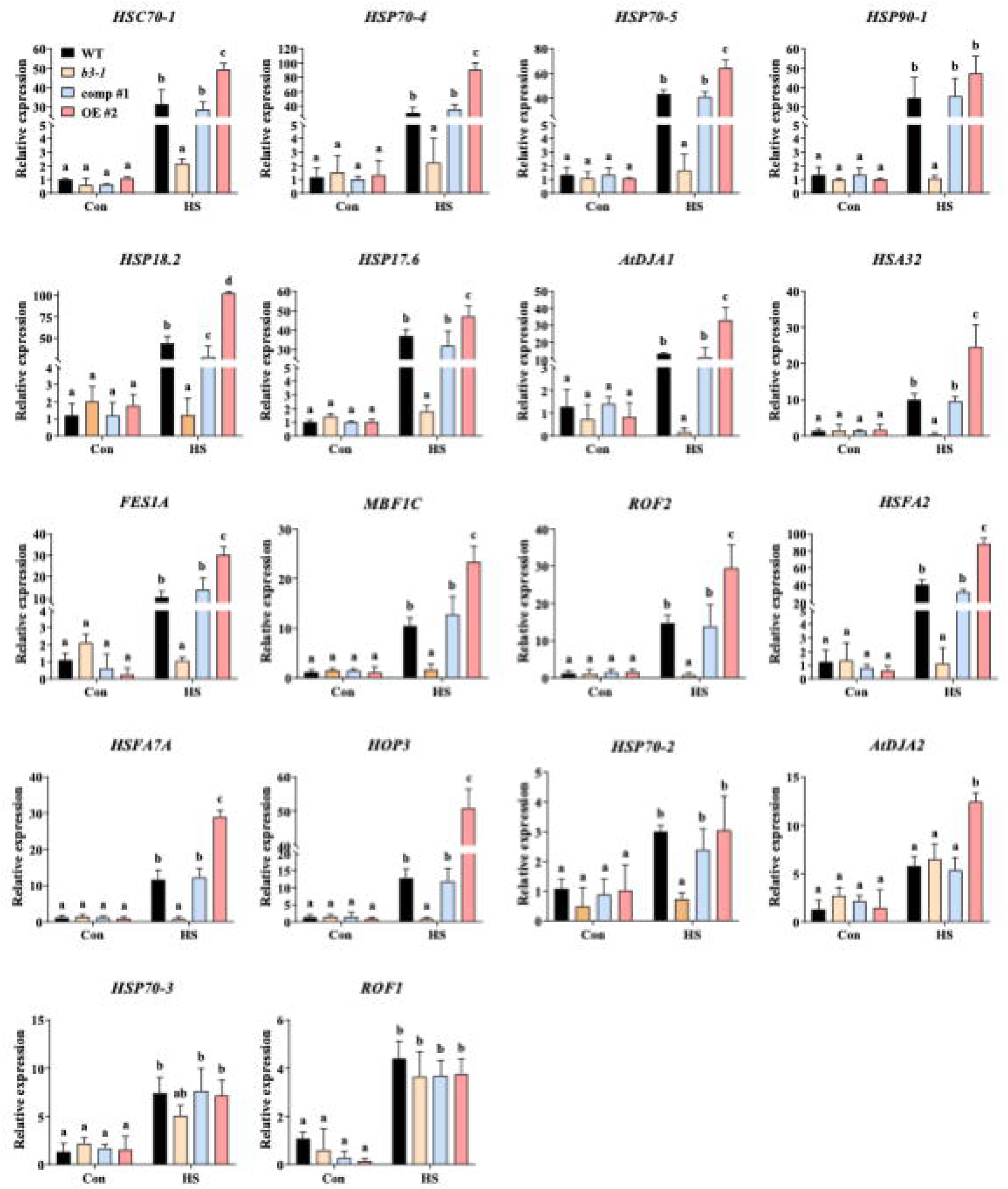
*AtDJB3* is required for the induction of heat-responsive genes during heat stress. Expression levels of several heat-responsive genes were analyzed using qRT-PCR in 1 O-day-old seedlings of WT, *AtDJB3* mutant *(b3-l),* complemented, and overexpression lines after exposure to 37°C for 2h. Transcript levels were normalized to *ACTIN2* and *GAPDH.* Data represent mean ± SE (w = 3). Statistical significance was determined using two-way ANOVA followed by Tukcy’s HSD test *(P <* 0.05); different lowercase letters indicate significantly different groups.

Among the other AtDjB paralogs studied in Arabidopsis, only AtDjB2 shares heat inducibility with AtDjB3. Despite functional redundancy observed in yeast complementation assays (Tak et al., 2023), *b2* mutant exhibited no transcriptional alterations upon heat treatment **(Supplementary** Fig. 4), suggesting AtDjB3 plays a predominant and more specific role in mediating transcriptional responses to high temperature.

### Loss of *AtDJB3* disrupts the targeting of Hsc70-1 to aggregated proteins under heat stress

Besides its pivotal role in solubilizing heat-induced protein aggregates, Hsp70 is also a central regulator of HSR, influencing Hsf activity under proteotoxic stress and ensuring cellular stability under adverse conditions (Masser et al., 2020; Mayer and Bukau, 2005; Richter et al., 2010). However, Hsp70s rely on JDP co-chaperones for all their functions (Kampinga and Craig, 2010). Our prior work showed that AtDjB3 associates with heat-induced protein aggregates and co-localizes with Hsp101, the major disaggregase for heat-denatured protein aggregates in Arabidopsis protoplasts, implicating AtDjB3 in aggregate clearance. Consistent with this, solubilization of heat-induced protein aggregates was severely compromised in *b3-1* mutant seedlings (Tak et al., 2023). o investigate the role of AtDjB3 in tethering Hsp70 to these heat-induced protein aggregates, we examined the subcellular localization of Hsc70-1-RFP, a cytosolic Hsp70 isoform expressed under its native promoter, under control and stress conditions in protoplasts of wild-type, *b3-1* and *AtDJB3* overexpression lines. Under control conditions, Hsc70-1 displayed a diffuse cytosolic distribution in all genotypes analyzed. However, following a treatment of 37°C, 30 min, Hsc70-1 rapidly transitioned to distinct punctae **(Fig. 4A)**, indicating its recruitment to misfolded protein aggregates. Quantitative analysis revealed an ∼85% reduction in Hsc70-1 puncta in *b3-1* protoplasts compared to WT, whereas protoplasts overexpressing AtDjB3 showed a marked increase in puncta formation **(Fig. 4B)**, positioning AtDjB3 as a crucial mediator in facilitating Hsc70-1 recruitment to misfolded proteins during heat stress.

**Figure 4.**
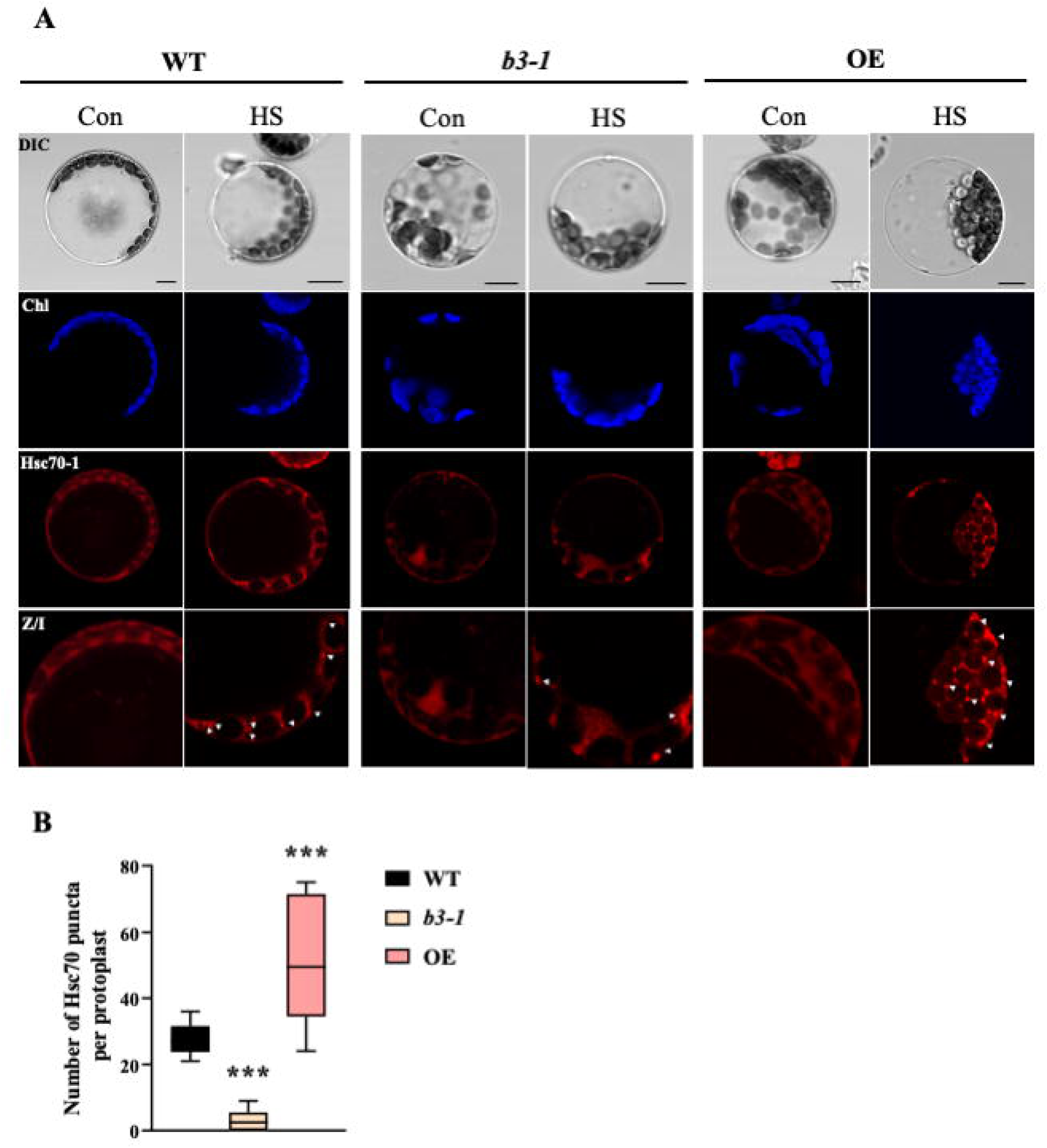
Loss of AtDjB3 decreases Hsc70-l puncta formation during heat stress. (A) Protoplasts were isolated >m four-week-old WT, *b3-l,* and OE seedlings and transiently transformed with the *pHSC70::HSC70-RFP* nstruct. Red fluorescence indicates Hsc70-l-RFP. while blue denotes chlorophyll autofluorcsccncc, visualized ing mRFP (572 nm/606 nm) and Cy5 (678 nm‘694 nm) filter sets, respectively. White arrows mark cytoplasmic >c70-l foci. (B) Quantification of Hsc70-l puncta per protoplast following 30 min heat stress at 37°C. Data were st tested for normality using the Shapiro-W’ilk test and for equal variance, prior to statistical analysis by one-way MOVA followed by Dunnctt’s post hoc test, with WT as the reference group (***, *P* < 0.0001). Box plots show the 11 range (minimum to maximum), interquartile range (Q1 —Q3), and median; horizontal lines represent mean ± SE *(n* 3; *N-* 4 protoplasts). Scale bar, 10 pm.

### AtDjB3 is essential for Hsc70-1 association with heat-induced misfolded proteins

To determine whether the reduction of Hsc70-1 localization to heat-induced protein aggregates in *b3-1* was due to altered total Hsc70-1 levels or its impaired recruitment to protein aggregates, total proteins isolated from WT, *b3-1*, and AtDjB3 overexpression lines following heat stress (35°C, 2 h) were fractionated into soluble and insoluble pools. Under non-stress conditions, Hsp70 distribution was comparable across all genotypes in both fractions **(Fig. 5A, left),** suggesting that the overall Hsc70-1 levels are not altered. However, its levels were markedly reduced in the insoluble fraction of *b3-1* after stress**(Fig. 5A, right)**. In contrast, *AtDJB3* overexpression enhanced Hsc70-1 accumulation in the insoluble fraction (Fig. 5B). Given the antibody’s cross-reactivity with other AtDjB paralogs (Tak et al., 2023), we validated AtDjB3 association using a transgenic line subjected to the same heat stress conditions. This confirms its presence in the insoluble fraction, indicating its association with heat-induced aggregates confirming its presence in heat-induced aggregates **(Supplementary** Fig. 5). These findings suggest that AtDjB3 promotes Hsc70- 1 recruitment to heat-induced aggregates in Arabidopsis.

**Figure 5.**
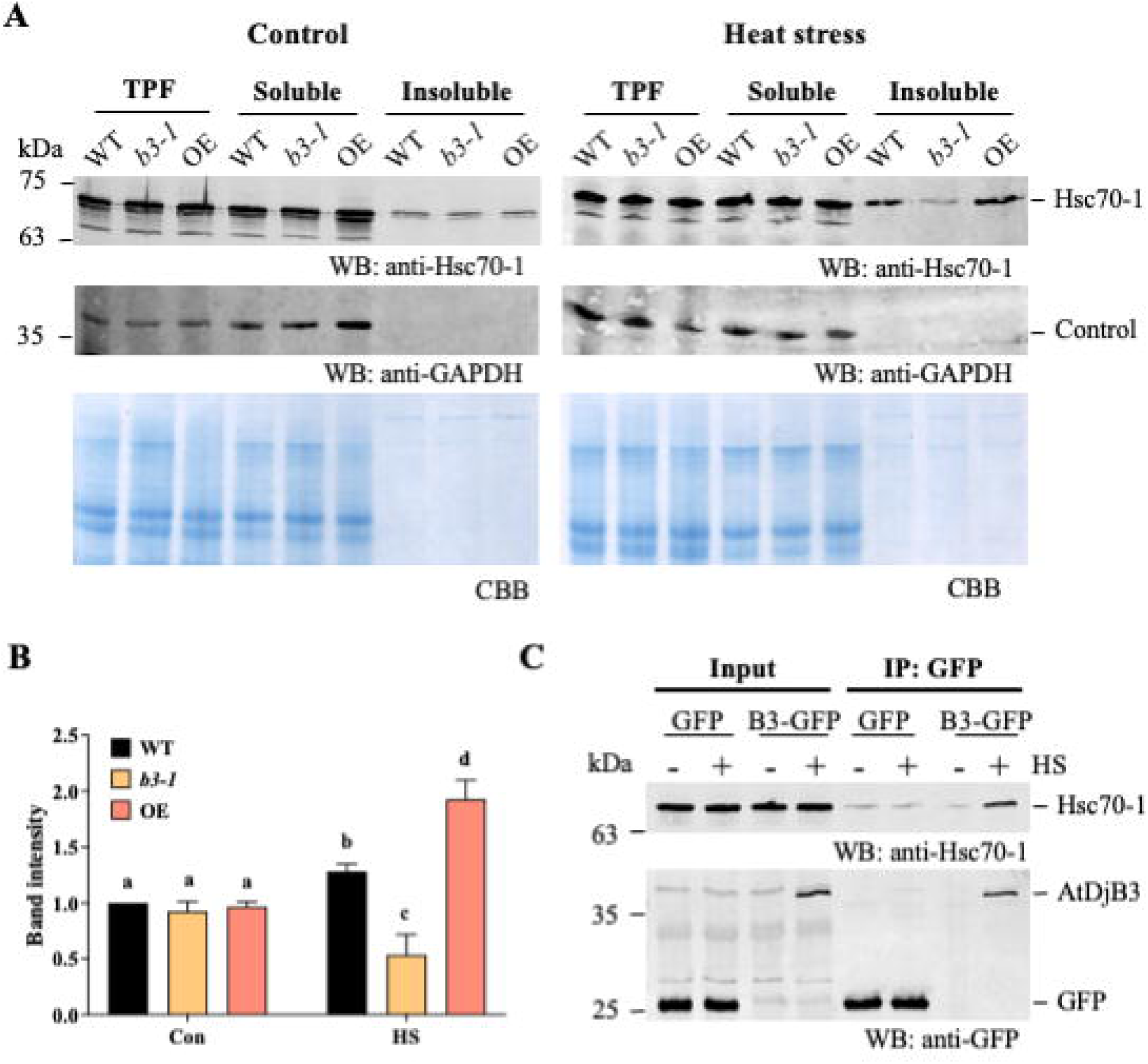
Loss of AtDjB3 leads to reduced association of Hsc70-l with protein aggregates. (A) 1 O-day-old WT. *b3-l, IDJB3* ovcrcxprcssion seedlings were exposed to heat stress at 35°C for 2h. Soluble and insoluble protein fractions separated by SDS-PAGE and probed with anti-Hsc70-l and anti-GAPDH antibodies. Coomassic Brilliant Blue ig was used to assess loading consistency. A representative blot of three independent experiments is shown. (B) tomctric quantification of Hsc70-l levels in the insoluble fraction, normalized to Coomassic staining. Statistical icancc was determined by two-way ANOVA followed by Tukcy’s HSD test; different lowercase letters denote icantly different groups (P < 0.05). Bars represent the mean ± SE of three biological replicates. (C) Co- noprccipitation of AtDjB3 and Hsc70-l was performed under both control and heat stress (35°C, 2h) conditions using *GFP* and *pAtDJB3B3::AtDJB3-GFP* transgenic lines. GFP-taggcd proteins were immunoprecipitated using GFP-Trap sc beads, and the eluates were probed with anti-GFP and anti-Hsc70-l antibodies to assess protein interactions.

### AtDjB3 associates with Hsc70-1 under heat stress *in vivo*

We next sought to determine whether AtDjB3 physically interacts with Hsp70 during heat stress. Co-immunoprecipitation (Co-IP) assays were performed using extracts from the control and heat-treated *AtDjB3*-GFP line described above. 35S::GFP line served as a constitutive GFP control. Upon heat exposure, Hsp70 was specifically enriched in the AtDjB3-GFP immunoprecipitates **(Fig. 5C)**, indicating a stress-dependent interaction. No such association was detected in the GFP-only samples, confirming interaction specificity. Put together, these results furnish a mechanistic explanation for AtDjB3-mediated recruitment of Hsc70-1 to heat-induced protein aggregates.

### AtDjB3 influences the nuclear localization of HsfA1d under heat stress

Hsc70-1 interacts with HsfA1d and HsfA1e, and attenuates HSR in Arabidopsis (Tiwari et al., 2020). Since AtDjB3 affected the heat-inducibility of several genes as well as the recruitment of Hsc70-1 to heat-induced protein aggregates, we set out to explore the molecular bases through which AtDjB3 influences HSR. For this, we first checked if AtDjB3 directly interacted with Hsfs. A yeast two-hybrid (Y2H) screen using a subset of Hsfs known to drive heat-responsive gene expression, including HsfA1a, HsfA1b, HsfA1d, HsfA1e, HsfA2, and HsfA7a was performed (Liu and Charng, 2013; Nishizawa-Yokoi et al., 2011; Schramm et al., 2006; Tiwari et al., 2020; Yoshida et al., 2011). Surprisingly, we could not detect a positive interaction between AtDjB3 and any of these Hsfs **(Supplemental Fig. 6)**.

Absence of a direct physical interaction between AtDjB3 and the Hsfs prompted us to consider alternative mechanisms through which AtDjB3 might influence HSR. Given its role in modulating Hsc70-1 activity, we hypothesized that AtDjB3 might be regulating HSR, by targeting the Hsc70-1-mediated attenuation of HsfA1d, a key transcription factor in heat-responsive gene expression (Tiwari et al., 2020; Yoshida et al., 2011). To test this, we examined the subcellular localization of HsfA1d in *b3-1* mutant under control and heat stress conditions using the differential centrifugation method described previously (Kubota et al., 2022). Under non-stress conditions, HsfA1d was predominantly cytosolic in all the genotypes analysed **(Fig. 6A, top)**. Following heat stress (35°C for 2h), HsfA1d accumulated in the nucleus in WT and AtDjB3 overexpression lines, consistent with its transcriptional activation role. In contrast, *b3-1* mutants exhibited markedly reduced nuclear localization of HsfA1d **(Fig. 6A, middle, bottom)**, indicating that AtDjB3 is required for the nuclear translocation of HsfA1d. To directly probe the regulatory mechanism, we performed co-immunoprecipitation to pull down Hsc70-1 using HsfA1d antibody from cytosolic protein extracts. Compared to WT and overexpression lines, subtle, but reproducibly higher levels of Hsc70-1 were pulled down in *b3-1* mutant. Densitometric analysis of Hsc70-1/HsfA1d ratios supports the idea that in *b3-1*, a statistically significant amount of Hsc70-1 remained associated with HsfA1d even during stress **(Fig. 6B)**. No signal was detected with control IgG **(Supplemental Fig. 7)**, confirming specificity. These findings suggest that AtDjB3 plays a role in the dissociation of HsfA1d from Hsc70-1- mediated repression, thereby enabling its nuclear entry and transcriptional activation.

**Figure 6.**
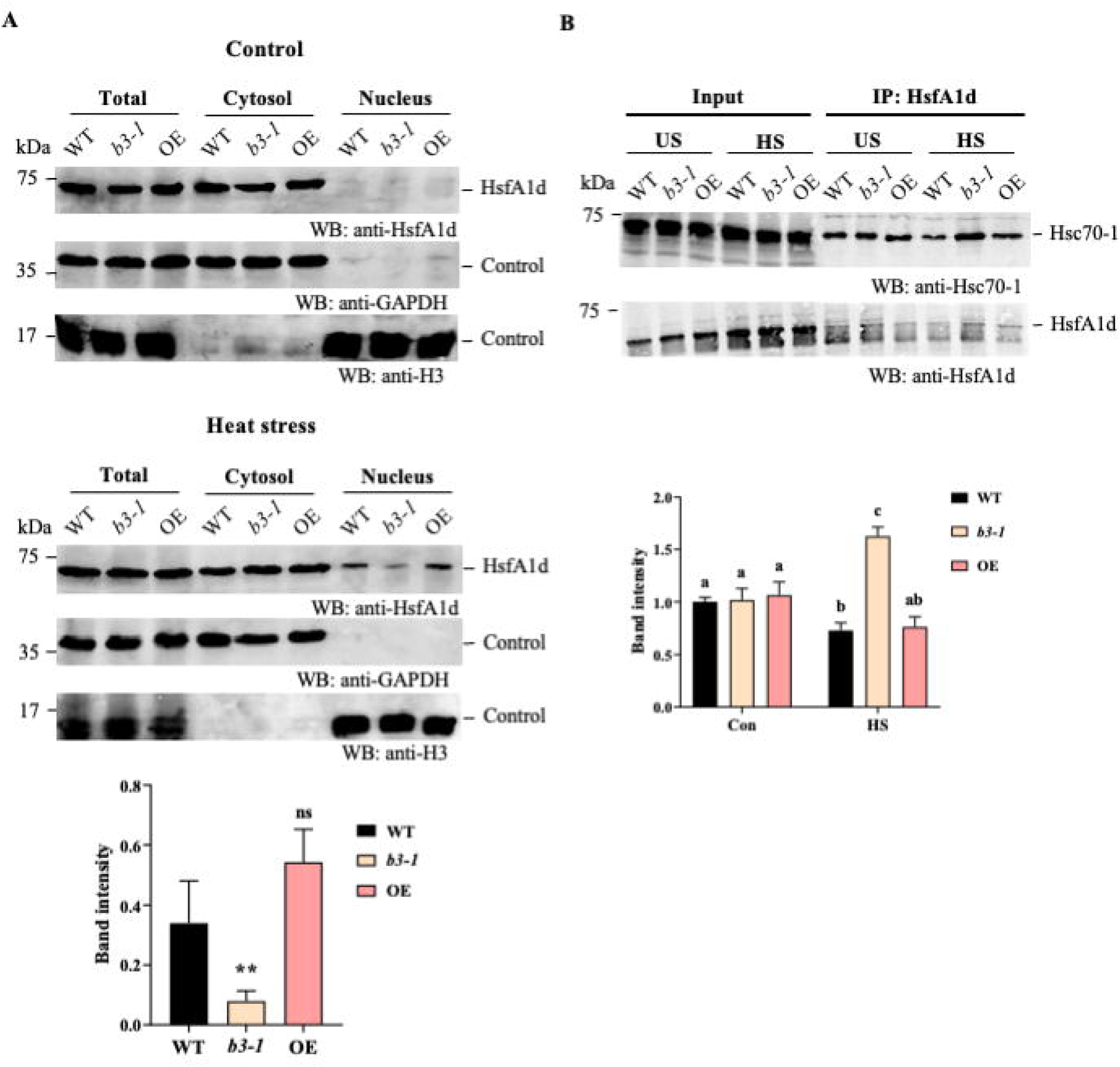
Loss of AtDjB3 impairs the nuclear localization of HsfAld under heat stress. (A) Nuclear and cytoplasmic onation was performed on 1 O-day-old seedlings of the indicated genotypes under control conditions (top) or after stress (35°C for 2h) (middle). Proteins from each fraction were immunoblottcd with anti-HsfAld antibody; anti- DH and anti-Histonc H3 antibodies were used as loading controls for the cytosolic and nuclear fractions, respectively, ircscntative blot of three independent experiments is shown. Nuclear HsfAld levels under heat stress were quantified vc to Histone H3 (bottom). Statistical analysis: one-way ANOVA with Dunnctt’s post hoc test (P < 0.05; *n* = 3). (B) Id was immunoprecipitated from total protein extracts of seedlings exposed to control or heat stress (35°C, 2 h). 0-1 co-immunoprccipitation was detected with anti-Hsc70-i antibody and normalized to HsfAld levels. Statistical sis: two-way ANOVA with Tukcy’s HSD test; different lowercase letters indicate significant differences (P < 0.05; *n = 3).*

## Discussion

Cells constantly manage cytotoxic protein aggregates through the coordinated action of the Hsp70:JDP-Hsp100 bi-chaperone system (Liberek et al., 2008). Additionally, Hsp70s sense the accumulation of these aggregates to induce an evolutionarily conserved heat shock response (HSR) that leads to the expression of Hsps to mitigate stress-induced damage and promote cell survival during or post-stress regimes (Masser et al., 2019; Mayer and Bukau, 2005). The present study provides evidence that AtDjB3, a JDP, plays a key role in modulating HSR by facilitating the recruitment of Hsp70 to heat-induced insoluble protein aggregates in Arabidopsis. This function promotes the nuclear localization of HsfA1d and, consequently, the expression of heat-inducible genes, thereby affecting thermotolerance.

Class II JDPs participate in myriad protein aggregate remodeling functions, including terminally misfolded proteins and amyloid aggregates across various organisms (Higurashi et al., 2008; Kirstein et al., 2017; Lopez et al., 2003; Wyszkowski et al., 2021). Previous studies have shown that AtDjB3 aids in the solubilization of heat-induced protein aggregates after heat shock, and its loss results in impaired acquired thermotolerance (Tak et al., 2023). However, a rapid and transient upregulation of AtDjB3 during heat stress, coupled with the basal thermotolerance defects observed in *AtDJB3* mutants, points to a role beyond aggregate remodeling. The fact that loss of AtDjB3 disrupted the heat-inducibility of several key heat-responsive genes, including *HSC70-1*, *HSP90-1*, *HSFA2*, *FES1A,* as well as small Hsps, strongly implicated AtDjB3 in the early stages of the heat-shock response in Arabidopsis.

Numerous studies have established that Hsp70 acts as a negative regulator of HSR (Craig and Gross, 1991; Lee and Schöffl, 1996). Hsp70 directly associates with Hsfs, repressing their activation under non-stress conditions. The general model posits that upon stress, Hsp70s are sequestered to misfolded protein aggregates, allowing Hsfs to translocate into the nucleus and activate heat-responsive genes (Bakery et al., 2024; Wang et al., 2023). However, not much is known about the role of molecular chaperones in HSR regulation of plants. Our study shows that in Arabidopsis, AtDjB3, a JDP co-chaperone of Hsp70, is critical for the efficient recruitment of Hsc70-1 to heat-induced aggregates during heat stress. This redistribution is crucial as it prevents Hsc70-1 from interacting with HsfA1d, a key transcriptional regulator, essential for the expression of many heat-responsive genes, including *MBF1C*, *HSP90*, *HSP18.2*, *HSFA2*, *HOP3*, *FES1A*, and *HSFA7A* (Kotak et al., 2007; Nishizawa-Yokoi et al., 2011; Yoshida et al., 2011). The fact that deletion of AtDjB3 impaired the nuclear localization of HsfA1d and disrupted the activation of these genes highlights its significance in modulating HsfA1d-dependent transcription. This function aligns with findings in other organisms, where JDP orthologs of AtDjB3 have also been implicated in transcriptional regulation under heat stress (Garde et al., 2023; Kmiecik et al., 2020; Marchler and Wu, 2001). In Drosophila and humans, DroJ1 and DnaJB1 attenuate the HSR by directly interacting with Hsf1’s transactivation domain, thereby preventing the recruitment of the transcriptional machinery (Kmiecik et al., 2020; Marchler and Wu, 2001; Shi et al., 1998). In contrast, AtDjB3 possibly operates much like Sis1 in budding yeast, which relocalizes to cytosolic foci upon heat stress, leading to the release of Hsf1 from Hsp70 and subsequent activation of the HSR (Ali et al., 2023; Feder et al., 2020).

Although all analyzed AtDjB proteins associate with heat-induced aggregates in Arabidopsis protoplasts, their specific contributions to the heat shock response (HSR) remain largely undefined. Notably, among these JDPs, only AtDjB2 and AtDjB3 are heat-inducible (Tak et al., 2023), yet AtDjB3 alone was found to influence transcriptional regulation during heat stress. This observation underscores the functional divergence among AtDjBs in their capacity to modulate the HSR in Arabidopsis. Class II JDPs have been implicated in both the activation and attenuation of HSR in other systems (Feder et al., 2020; Kmiecik et al., 2020; Marchler and Wu, 2001), suggesting that further comprehensive studies are needed to determine whether other AtDjBs play regulatory roles in HSR.

Interestingly, while our study primarily focused on the regulation of HsfA1d, the complexity of the Hsf network in *Arabidopsis thaliana*, along with evidence that Hsc70-1 also directly interacts with HsfA1e (Tiwari et al., 2020), raises the possibility that AtDjB3 may function beyond the Hsc70-1:HsfA1d module. Interestingly, a recent study reported that HsfA1d contains prion-like domains (PrDs) capable of undergoing heat-induced phase separation to form transcriptionally active condensates, which enhance gene expression (Peng et al., 2025). By titrating Hsc70-1 away from HsfA1d, AtDjB3 may promote the formation of these condensates and amplify Hsf-driven transcriptional activation. Although our yeast two-hybrid assay did not detect a direct interaction between AtDjB3 and HsfA1d, it remains possible that other AtDjBs interact with Hsfs containing prion-like domains, particularly given the known ability of AtDjBs to remodel prion-like aggregates in yeast (Tak et al., 2023). These findings point to a potential broader role of JDPs in modulating the phase behavior and activity of heat shock transcription factors in plants.

In conclusion, this study identifies AtDjB3 as a central regulator of the heat shock response (HSR) in Arabidopsi*s*, linking protein quality control with transcriptional regulation. By promoting the recruitment of Hsc70-1 to heat-induced protein aggregates, AtDjB3 supports proteostasis while also facilitating the nuclear localization of HsfA1d. This dual role reveals a mechanistic link between chaperone activity and stress-responsive transcriptional activation **(Fig. 7)**. Overall, our findings highlight the critical interplay between chaperone-mediated proteostasis and Hsf-regulated gene expression in plant thermotolerance, emphasizing the broader importance of J-domain proteins in plant stress resilience.

**Figure 7.**
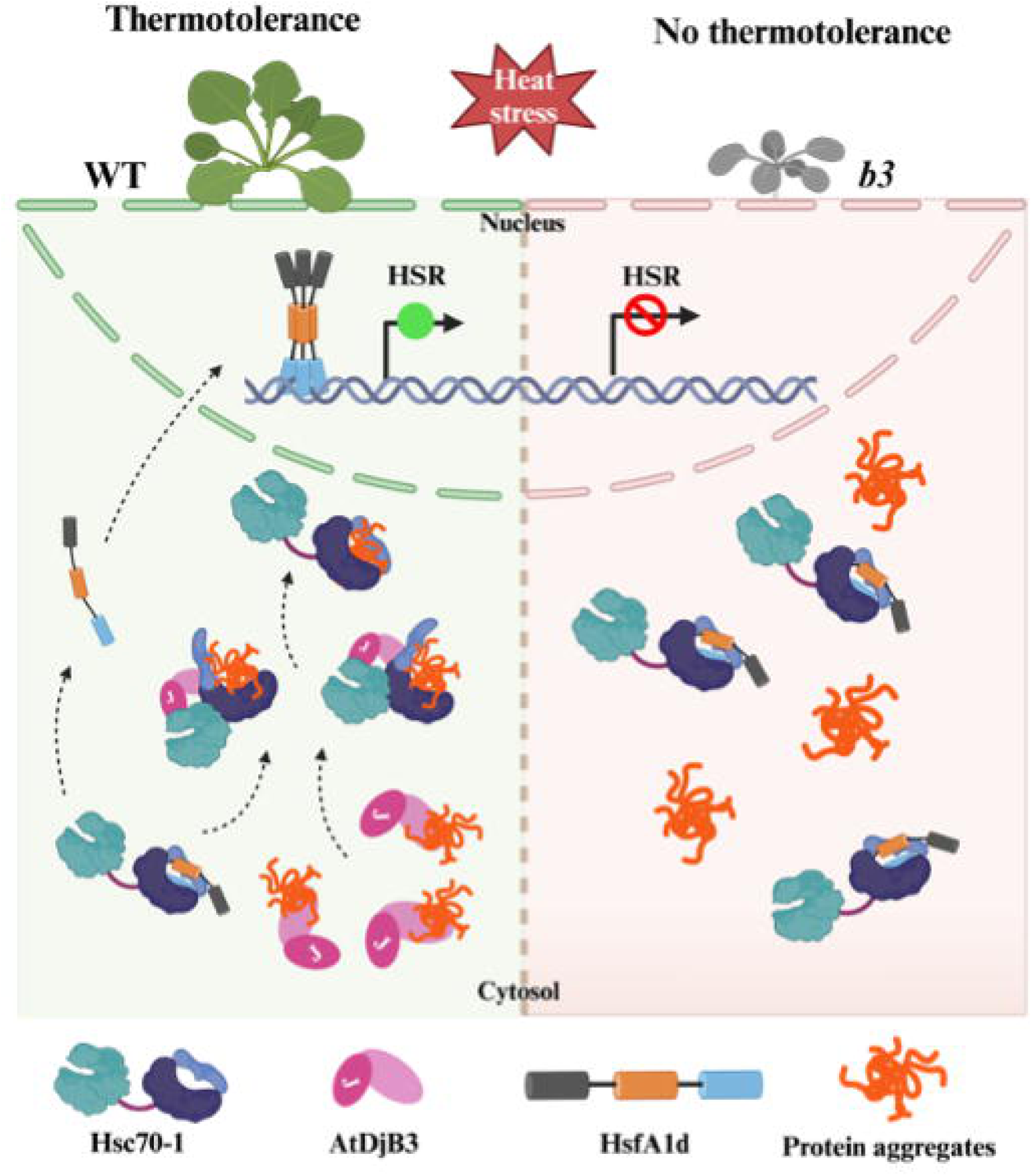
AtDjB3 functions in regulating HSR in Arabidopsis. Under heat stress conditions, AtDjB3 identifies protein aggregates formed due to elevated temperatures. This aids in the recruitment of Hsc70-l to aggregated proteins, thereby freeing HsfAld. HsfAld then translocates into the nucleus, where it trimcrizcs and activates HSR. ultimately promoting thcrmotolcrancc (left panel). In the absence of AtDjB3, Hsc70-l remains associated with HsfAld in the cytosol, thereby attenuating HSR. resulting in thermosensitivity (right panel).

## Materials and Methods

### Plant materials and growth conditions

*Arabidopsis thaliana* ecotype Col-0 was used as the wild-type background, as well as for mutant and overexpression lines, in all experiments. Surface-sterilized seeds were plated on ½ MS medium containing agar (0.7%) and sucrose (1%). These plates were placed for 48h at 4°C for stratification and then transferred to a Percival LED22C8 growth cabinet at 22 °C day/18 °C night and 70% humidity (light intensity 120 ± 20 μmol m^−2^ s^−1^, 16 h light:8 h dark cycle). The T-DNA insertion lines of *AtDJB3* (*b3-1*; SALK_003982 (already characterized), *b3-2*; SALK_051100C) were obtained from the Arabidopsis Biological Resource Center (ABRC; https://abrc.osu.edu). The mutant line was screened for homozygosity by genomic PCR and qRT-PCR using primers specified in **Table S1**.

GFP, overexpression, and complemented transgenic lines of AtDjB3 were generated using Agrobacterium tumefaciens (strain GV3101)-mediated floral-dip transformation. Positive transformants for AtDjB3-GFP were identified by germinating T0 seeds directly in the soil, followed by selection through spraying with 0.1% BASTA herbicide. In contrast, overexpression and complemented lines were screened by plating the seeds on a hygromycin-containing (50 mg/ mL) selection medium. The expression of the transgene in selected lines was further confirmed by qRT-PCR.

### Plasmid constructions

To generate *pHSC70::HSC70-RFP* for subcellular localization studies, an 1884bp promoter of *HSC70-1* was amplified from the genomic DNA of wild-type plants, and its full-length CDS of *HSC70-1* was sewn together using overlapping PCR. The resulting fragment subsequently shuttled into the pGWB653 destination vector with a C-terminal RFP tag. Similarly, the *pAtDJB3::AtDJB3-GFP* construct was created by amplifying a 1018 bp *AtDJB3* promoter from wild-type genomic DNA, sewing together the full-length CDS using overlapping PCR, and inserting it into pGWB604 (Nakagawa et al., 2007) to drive expression under its native promoter. To generate overexpression lines, the full-length CDS of *AtDJB3* was cloned into the HBP047 vector under the control of the constitutive CaMV 35S promoter. For the Yeast Two-Hybrid (Y2H) assay, the full-length CDS of *AtDJB3* was cloned into the pGADT7 destination vector, while the full-length CDS of *HSFA1A*, *HSFA1B*, *HSFA1D*, *HSFA1E*, *HSFA2*, and *HSFA7A* were cloned into the pGBKT7 destination vector to investigate their interactions. For protein expression and purification, the full-length CDS of *HSC70-1* and *HSFA1D* were cloned into the BamHI-SalI sites of the pSMT3 vector (kind gift from Dr. Christopher D Lima, USA) with an N-terminal His+SUMO tag.

### Seedling thermotolerance assay

Surface-sterilized seeds from wild type, *AtDJB3* mutants, complemented, and overexpression lines were plated on ½ MS media and grown for 4 days (Percival LED22C8 growth chamber; 22 °C day/18 °C night). Seedlings with expanded green cotyledons were heat-stressed at 37°C under long-day light conditions for 2 days, followed by a 12-day recovery (22 °C day/18 °C night, 16h light/8h dark, 120±20 μmol m^−2^ s^−1^). Survival was assessed by counting seedlings with open green cotyledons and documented through photography. Each genotype included 150 seedlings, and trials were repeated at least four times.

### Fresh weight measurement

Seedlings were collected before and after heat treatment, rinsed with distilled water, blotted dry, and weighed. Two batches were measured per experiment, and the procedure was repeated three times. The average weight and standard deviation were calculated.

### Chlorophyll estimation

Chlorophyll estimation was performed as mentioned previously (Tak et al., 2023) some modifications. 150 seedlings per experiment were incubated in 95% ethanol at 4°C in the dark for 20h with gentle rotation. After centrifugation (10,000 g for 10 min, 4°C), the supernatant was analyzed spectrophotometrically at 664 nm and 649 nm. Chlorophyll a (Chl a) and chlorophyll b (Chl b) concentrations were calculated using: Chl a (mg/L) = 13.36 A664 – 5.19 A649; Chl b (mg/L) = 27.43 A649 – 8.12 A664.

Total Chlorophyll (mg/L) = Chl a + Chl b

To calculate chlorophyll content in terms of plant tissue:

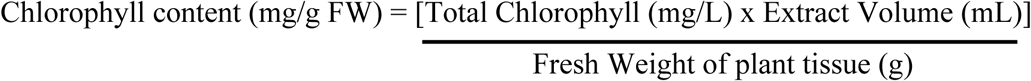

### Protoplast-based transient expression analysis

The protoplast isolation protocol followed in the study was used as described previously (Yoo et al., 2007). Briefly, the protoplasts of WT, *b3-1,* and *AtDJB3* overexpression were transformed with *pHSC70::HSC70-1-RFP* and were observed before and after HS under a live-cell microscope (Olympus FluoView-3000). Excitation and emission wavelengths Ex488/Em500-520 and Ex594/Em630-700 were used for GFP and chlorophyll, respectively. The images acquired from the confocal microscope were analyzed using ImageJ software.

### Yeast two-hybrid assays

The protein interaction study was performed as described previously (Tamadaddi *et al*., 2021). Briefly, the respective combinations of AtDjB3 and each Hsfs were co-transformed into the Y2Hgold yeast strain. Yeast transformants were initially selected on SD/-Leu-Trp double dropout plates. The interaction was further confirmed by growing on the selection medium SD/-Leu-Trp-His triple dropout plates.

### RNA isolation and qRT-PCR

Total RNA was isolated from 10-day-old seedling tissues under control and heat stress conditions as mentioned previously (Tak et al., 2023). cDNA was prepared using an iScript cDNA synthesis kit (Bio-Rad). Real-time PCR was performed in a CFX Connect 96-well real-time system using the Green Eye Ab universal qPCR Master Mix (Genes2me). Three biological replicates were analyzed for each sample with their respective technical replicates. Reference genes used were *ACTIN2* and *GAPDH* for normalization. Sequences of all the primers used are listed in **Table S1**.

### Isolation of soluble and insoluble protein fractions from Arabidopsis seedlings

Protein fractionation into soluble and heat-denatured components was performed as described previously (Tamadaddi *et al*., 2021). Briefly, 10-day-old Arabidopsis seedlings grown on ½ MS plates were harvested either as untreated controls or following heat stress at 35°C for 2h. For each sample, 0.7 g of tissue was ground in protein isolation buffer (25 mM HEPES, pH 7.5, 200 mM NaCl, 0.5 mM EDTA, 0.1% Triton X-100, 5 mM ε-amino-N-caproic acid, 1 mM benzamidine-HCl) and homogenized using a Dounce grinder on ice. The crude extract was centrifuged at 16,100 g for 15 min to separate soluble and insoluble fractions. The soluble fraction was collected, while the pellet was washed with quartz salt and protein isolation buffer (with and without Triton X-100) before resuspension in the SDS-PAGE sample buffer. Both fractions were analyzed via SDS-PAGE and immunoblotting.

### Arabidopsis protein extraction and fractionation

Protein extraction from the cytosolic and nuclear fractions of Arabidopsis was conducted with modifications to an established protocol (Kubota et al., 2022). 10-day-old seedlings (1.5g), either untreated or heat stress-treated, were ground in liquid nitrogen. Crude protein extracts were obtained by homogenizing the tissue in 2 mL of NIB buffer (20 mM Tris-HCl, 20 mM MgCl2, pH 6.8, 5% sucrose, 40% glycerol, 0.8% Triton X-100, 0.08% β- mercaptoethanol, protease inhibitor, 1 mM DTT, and 1.3 mM PMSF) followed by filtration through Miracloth. 300 μL of filtrate was mixed with 100 μL of SDS sample buffer to obtain the total protein fraction. For cytosolic proteins, 1 mL of filtrate was centrifuged at 12,000 g for 10 min at 4°C, and the supernatant was combined with 100 μL of SDS sample buffer. The pellet was washed six times with a NIB buffer containing Triton X-100, followed by a final wash with ice-cold NIB buffer without Triton X-100. The pellet, containing the nuclear protein fraction, was resuspended in NIB buffer (without Triton X-100), mixed with SDS sample buffer, and briefly centrifuged (1,500 g, 30 secs) to collect the supernatant.

### Co-immunoprecipitation (Co-IP)

Co-immunoprecipitation was performed using GFP-Trap® Agarose beads (ChromoTek) following the manufacturer’s protocol. 10-day-old transgenic seedlings expressing either GFP alone or AtDjB3-GFP were heat-stressed at 37°C for 2h. 1 g of seedlings was ground in liquid nitrogen and homogenized in extraction buffer (25 mM HEPES, pH 7.5; 100 mM NaCl; 10 mM MgCl₂; 10% glycerol; 5 mM DTT; 1% Triton X-100; protease inhibitors). After vortexing and incubation on ice for 30 min, the lysate was clarified by centrifugation (1,500 g, 30 sec). The supernatant was incubated with 20 μL of pre-equilibrated agarose beads for 45 min at 4°C, followed by three washes with a high-salt extraction buffer (400 mM NaCl). Bound proteins were eluted in 2× SDS sample buffer (120 mM Tris-HCl, pH 6.8; 20% glycerol; 4% SDS; 0.04% bromophenol blue; 10% β-mercaptoethanol) by heating at 95°C for 5 min and analyzed via immunoblotting.

To examine Hsp70-HsfA1d association, co-immunoprecipitation was performed on cytosolic extracts from 10-day-old WT, *b3-1*, and *AtDJB3* overexpression seedlings, either untreated or heat-stressed. Extracts were incubated overnight at 4°C with protein A magnetic beads (Invitrogen) pre-bound to an anti-HsfA1d antibody (in-house generated) or mouse IgG (negative control). After thorough washing, bound proteins were eluted with 2× SDS sample buffer at 95°C and analyzed by Western blot. As a negative control, a parallel immunoprecipitation was performed using non-immune mouse IgG under identical conditions to confirm the specificity of the interaction.

### Protein purification and antibody generation

A single colony of *E. coli* Rosetta cells, containing the target protein expression vector, was cultured at 37°C until OD600 reached 0.6-0.8. Protein expression was induced with 0.5 mM IPTG at 16°C overnight. Cells were harvested by centrifugation and resuspended in lysis buffer (50 mM Tris-HCl pH 7.5, 100 mM KCl, 20 mM imidazole, 0.5% Triton X-100, 10% glycerol, protease inhibitors, 20 mM β-mercaptoethanol, 1 mM DTT, 1 mM PMSF, 1 mg/ mL lysozyme). The suspension was sonicated and centrifuged. For protein binding, the clarified lysate was incubated with pre-equilibrated Ni-NTA agarose beads (Qiagen). Beads were washed with a lysis buffer to remove non-specific proteins. The target protein was eluted with an elution buffer (50 mM Tris-HCl, pH 7.5, 100 mM KCl, 250 mM imidazole, 10% glycerol). Purified Hsc70-1 and HsfA1d proteins were dialyzed and cleaved to remove the His+SUMO tag. Tag-free protein was used for antibody generation at our In-house animal facility.

### Immunoblot

Protein samples were resolved by SDS-PAGE and transferred onto a methanol-charged PVDF membrane using the wet transfer method. Primary antibodies used in the study: GAPC (1:3000; Agrisera AS15 2894), Histone H3 (1:3000; Agrisera AS10 710); Hsc70-1 and HsfA1d (in-house generated; 1:1000). Blots were then incubated with Invitrogen’s Goat anti-rabbit or anti-mouse (IgG (H+L) DyLight 680 or 800) secondary antibodies (1:20000 dilution) and visualized using a Li-Cor Odyssey system.

### Accession numbers

The sequence details for the genes covered in this study were obtained from The Arabidopsis Information Resource (TAIR) databases under the following accession numbers: *AtDJB3* (AT2G20560), *HSC70-1* (AT5G02500), *HSP70-2* (AT5G02490), *HSP70-3* (AT3G09440), *HSP70-4* (AT3G12580), *HSP70-5* (AT1G16030), *HSP90-1* (AT5G52640), *HSA32* (AT4G21320), *HSP18.2* (AT5G59720), *HSP17.6* (AT5G12030), *AtDJA1* (AT3G44110), *AtDJA2* (AT5G22060), *FES1A* (AT3G09350), *MBF1C* (AT3G24500), *ROF1* (AT3G25230), *ROF2* (AT5G48570), *DREB2A* (AT5G05410), *APX2* (AT3G09640), *HSFA1A* (AT4G17750), *HSFA1B* (AT5G16820), *HSFA1D* (AT1G32330), *HSFA1E* (AT3G02990), *HSFA2* (AT2G26150), *HSFA7A* (AT3G51910), *ACTIN2* (AT3G18780)*, GAPDH* (AT3G26650).

## Acknowledgements

The authors thank Surekha Katiyar Agarwal (DU, Delhi) for Hsf plasmids and Fund for Improvement of Science & Technology Infrastructure in Higher Educational Institutions (FIST) for the live-cell imaging system at IISER Bhopal. Figures were created with (BioRender.com (https://biorender.com).

## Author Contributions

G.S. and C.S. conceptualized the study, designed the experiments, and contributed to manuscript writing. C.S. supervised the research, secured funding, and facilitated laboratory resources. G.S. performed experiments and analyzed the data. I.S. contributed to resource generation and the Y2H assay.

## Data availability

The authors confirm that all experimental data are available in the main text and/or the supplementary materials.

