## Supplemental table for "AtDjB3 regulates Hsc70-1-mediated expression of the heat shock genes and thermotolerance in Arabidopsis"

### Supplementary figures and tables

**Supplementary table 1. (Table S1).** List of primers used in this study

| RT primer | | |
|---|---|---|
| *HSC70-1* | AAGGAAACAGAACCACGCCA | TGTCAGAGAAACGACGACCG |
| *HSP70-2* | CCTGATGAGGCTGTAGCTTATG | CTTCTTGGTTGGAATGGTTGTG |
| *HSP70-3* | TAACTTTCTCTCTTACTCTCTT | CAAAACATTCCTGTTATAGATTA |
| *HSP70-4* | GCCTTTTGGCTTTTGTTTACT | AACGGTAAAGCTAAATTCTTAAA |
| *HSP70-5* | AGCGTGTTTAGTGTGTTAAG | CTCGAAACAACTGAAACAGC |
| *HSP90-1* | CAACACGTTCTACAGCAACAAAG | AGCTCTCAAATCGGATCTTGTC |
| *HSP18.2* | GGATTCTTCACGCCATCTTCTG | CTCTCTCCGCTAATCTGCAG |
| *HSP17.6 II* | AGACATGCTTGAAGTCCCCG | ATGACGTCAGCAGGTGTAGC |
| *FES1A* | GGATCGAAAATGGTTTATGGAG | CGATCGATTCAACGTGCTC |
| *HSA32* | GGTGATTGGGCTGAACATATG | GGCTCTGAGACCACCGTTC |
| *MBF1C* | GACGATGCCGAGCAGATACC | TTTCGGATCGCGTAGGTCTT |
| *ROF1* | GCGAAACTAAGTAAGGAATAATCA | GCAACACATTGGGATAATCTC |
| *ROF2* | GATTCGAGCCGTGACAGAG | CCATGCTATCAACTCCACATC |
| *AtDJA2* | CCTTGTGTGGCTTCCAGTTT | CGTGAAGTGAATGTATAGCTTACCC |
| *AtDJA1* | AGCTTTGTGTGGCTTCCAAT | GAAGTGGATGTAGAGCTTTACCC |
| *AtDJB3* | GGCTGCTCCTATCGAAAACAAG | GGCATCCCTTCCTTCGGTAC |
| *HSFA7A* | CTCAAGTTTCATCAGACAAC | CTGCGAAGCTCGTTGCAAGC |
| *HOP3* | TTTATCTTGAGATGGGGAAG | GTATCTGGATTACGATGCTCTG |
| *HSFA2* | AACAGCTTTGTGGTGTGGGA | TGAATCCATAAGTATTGAGCTGACG |
| *APX2* | CTTGATGATCCTCTCTTTCTCCCA | ACTCCTTGTCAGCAAACCCGAG |
| *DREB2A* | CAGTGTTGCCAACGGTTCAT | AAACGGAGGTATTCCGTAGTTGAG |
| *ACTIN2* | TATCGCTGACCGTATGAGCAAAG | TGGACCTGCCTCATCATACTCG |
| *GAPDH* | TTGGTGACAACAGGTCAAGCA | AAACTTGTCGCTCAATGCAATC |

| | | |
|---|---|---|
| **Genotyping primers** | | |
| *AtDJB3*-SALK lines | CTCCTGTCTTCTGCTCTGAGG | TGCTGTTGCATATGGTAGCTG |
| **Gateway cloning primers** | | |
| *AtDJB3* (with stop codon) | GGGGACAAGTTTGTACAAAAAAGCAGGCTGGATGGGAGTGGATTACTACAAAG | GGGGACCACTTTGTACAAGAAAGCTGGGTGTCATCCGAGAAGTTTCTTCACTC |
| *pAtDJB3::AtDJB3-GFP* F1 | GGGGACAAGTTTGTACAAAAAAGCAGGCTGGGGAGATTGTAGACCTCTAGGTTTC | GTAGTAATCCACTCCCATCCGATCTTCTTCTCTCTTTCGC |
| *pAtDJB3::AtDJB3-GFP* F2 | GCGAAAGAGAGAAGAAGATCGGATGGGAGTGGATTACTAC | GGGGACCACTTTGTACAAGAAAGCTGGGTGTCCGAGAAGTTTCTTCACTCCTGT |
| *pHSC70::HSC70-RFP* F1 | GGGGACAAGTTTGTACAAAAAAGCAGGCTGGCGCGTTAGCTTTATCTGCATCC | GTCCTTCTCCTTTACCCGACATTTTTATCGGAAGATTTGGAAAC |
| *pHSC70::HSC70-RFP* F1 | GTTTCCAAATCTTCCGATAAAAATGTCGGGTAAAGGAGAAGGAC | GGGGACCACTTTGTACAAGAAAGCTGGGTG GTCGACCTCCTCGATCTTAGG |
| **Cloning primers** | | |
| HsfA1d | GTCAGGATCCATGGATGTGAGCAAAGTAACC | GACTGTCGACTCAAGGATTTTGCCTTGAGAG |
| Hsc70-1 | CGCGGATCCATGTCGGGTAAAGGAGAAGGACC | CCCAAGCTTTTAGTCGACCTCCTCGATCTTAGGTC |
