## Supplemental figures for "AtDjB3 regulates Hsc70-1-mediated expression of the heat shock genes and thermotolerance in Arabidopsis"

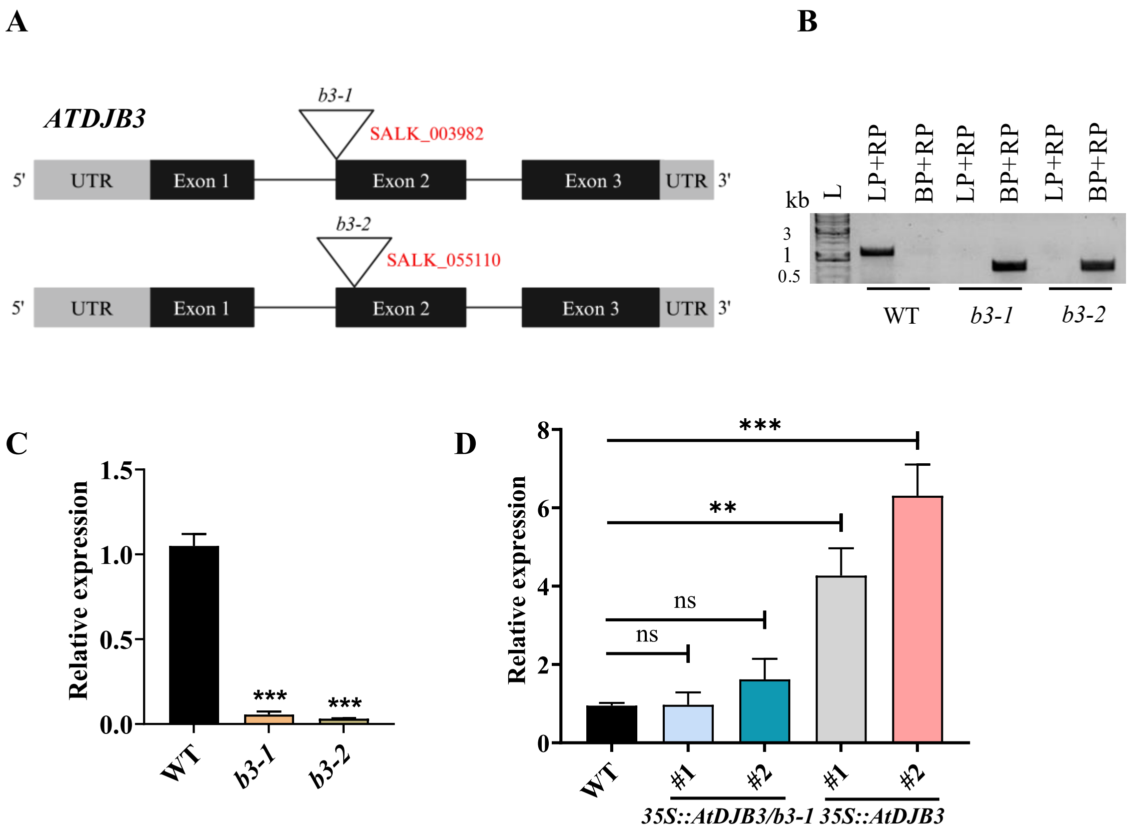


**Supplemental Figure 1. Characterization of T-DNA insertion mutants, complementation, and overexpression lines of *AtDJB3* in *Arabidopsis thaliana*. (A)** Schematic representation of the *AtDJB3* gene structure, where black boxes, black lines, and grey boxes correspond to exons, introns, and untranslated regions (UTRs), respectively. Inverted triangles mark the T-DNA insertion. **(B)** Genotyping PCR analysis confirming the presence of T-DNA insertions in *b3-1* and *b3-2* mutants. **(C)** qRT-PCR analysis of *AtDJB3* transcript levels in *b3-1* and *b3-2*, using primers designed to amplify regions downstream of the T-DNA insertion sites. **(D)** Relative *AtDJB3* mRNA expression levels in complemented and overexpression lines were determined via qRT-PCR. *ACTIN2* and *GAPDH* served as internal reference genes for normalization. Statistically significant differences in expression levels are denoted by asterisks (Dunnett's test, one-way ANOVA), while 'ns' indicates non-significant comparisons ($P > 0.05$).

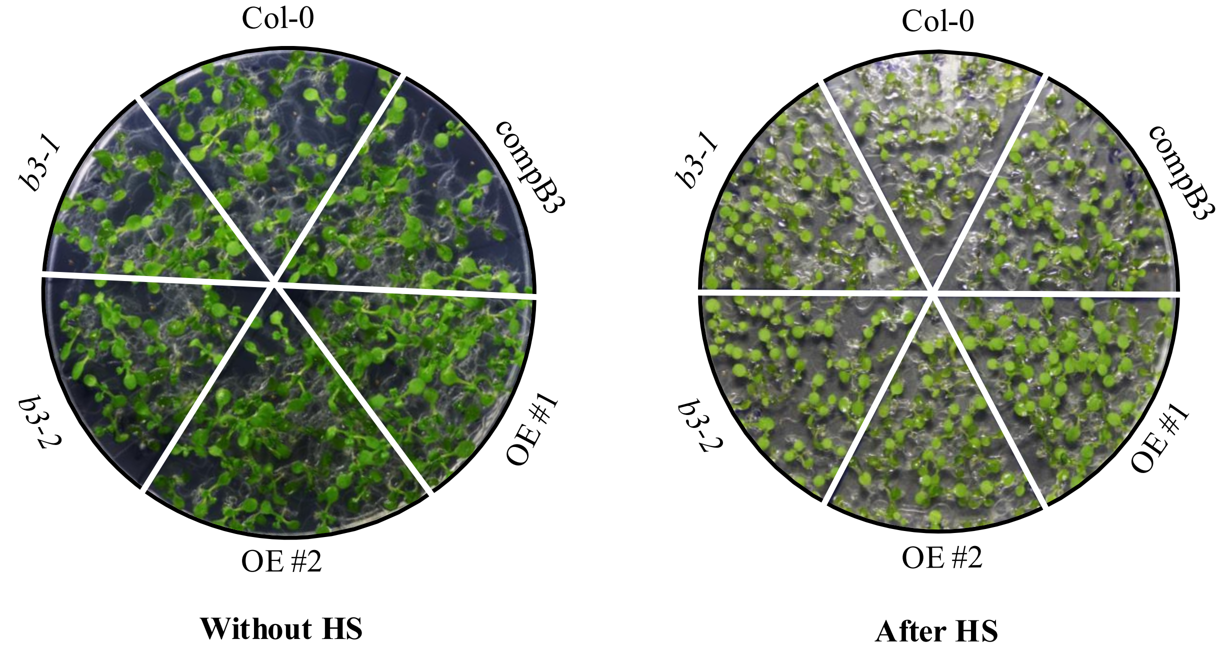


**Supplementary Figure 2. Thermotolerance phenotype of AtDjB3 lines following short-term heat stress.** 10-day-old seedlings of WT, *AtDJB3* mutants (*b3-1* and *b3-2*), complemented, and overexpression lines (OE #1 and #2) were subjected to heat stress at 37°C for 2h. Seedling growth and recovery were photographed on day 12 post-treatment.

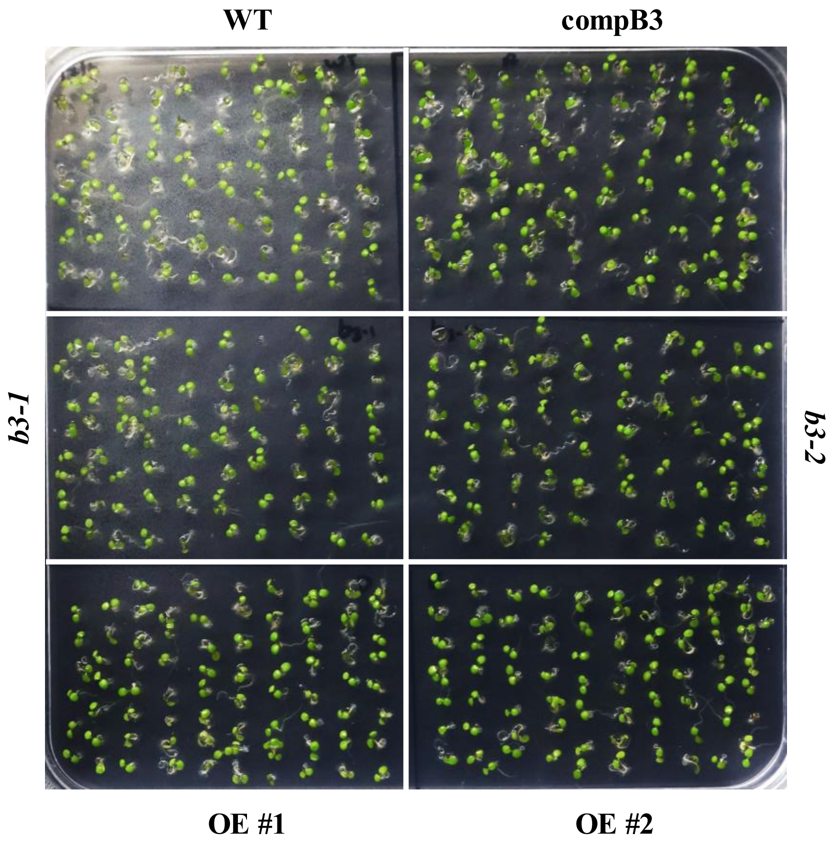


**Supplementary Figure 3. Basal growth phenotype of WT, AtDjB3 mutants, and overexpression lines under non-stress conditions.** Seedlings were assessed 4 days after germination under control conditions, prior to exposure to heat stress.

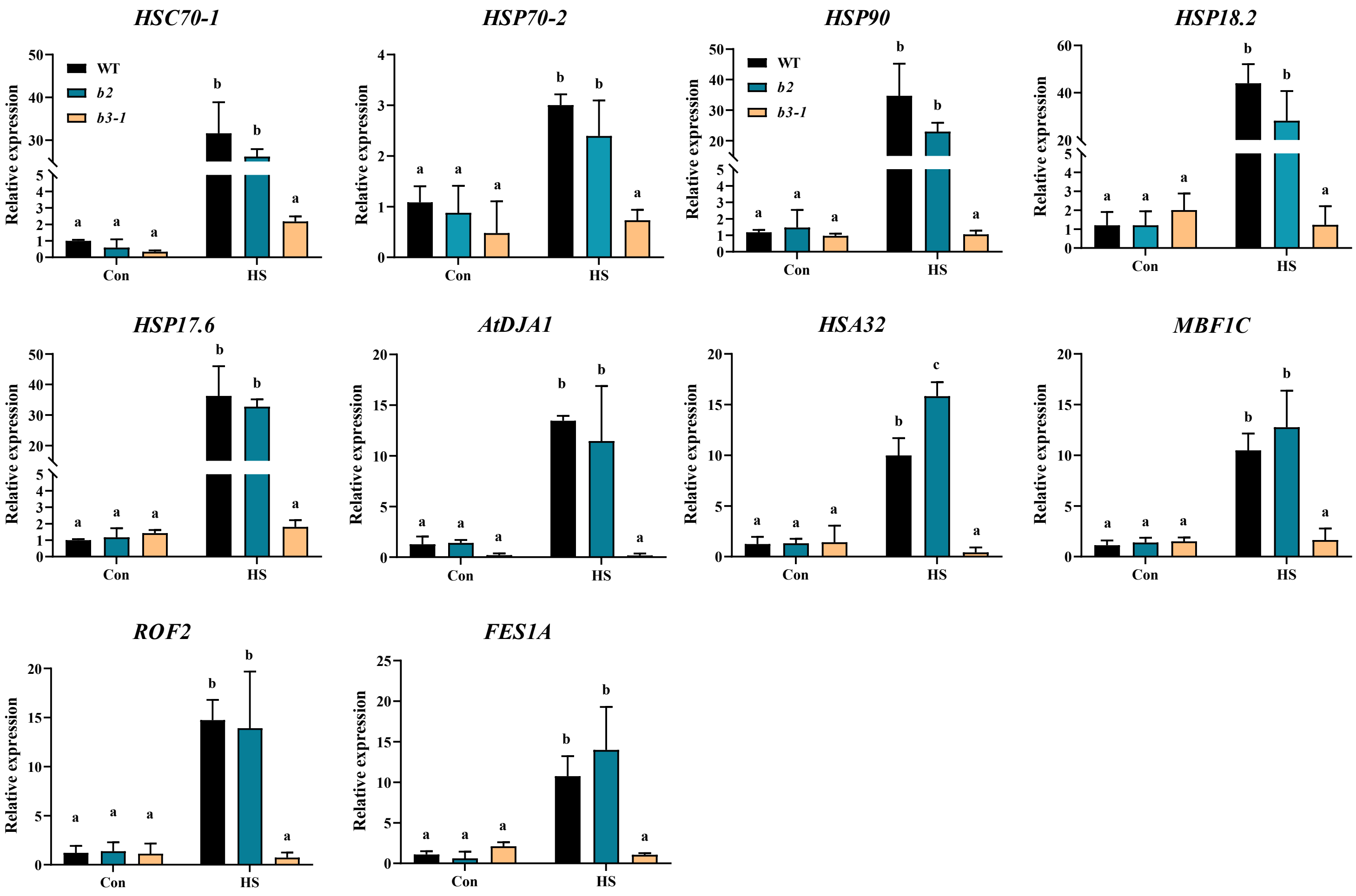


**Supplementary Figure 4. *AtDJB2* is not required for the expression of heat-responsive genes during heat stress.** qRT-PCR analysis of heat-responsive genes in 10-day-old seedlings of WT, *AtDJB2* (*b2*), and *AtDJB3* mutant (*b3-1*) lines following 2 h heat stress at 37°C. Transcript levels were normalized to *ACTIN2* and *GAPDH*. Statistical differences were assessed using two-way ANOVA followed by Tukey's HSD test ($P<0.05$); different letters above bars indicate significantly distinct groups.

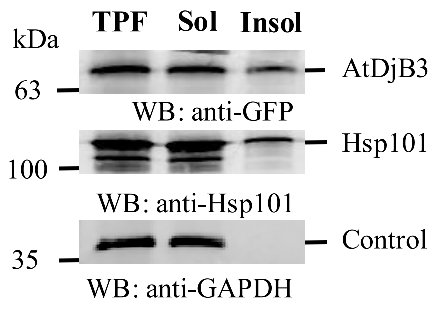


**Supplementary Figure 5. AtDjB3 associates with heat-induced aggregates.** 10-day-old *AtDjB3–GFP* seedlings were subjected to heat stress at 35 °C for 2h. Following treatment, soluble and insoluble protein fractions were isolated and subjected to SDS-PAGE. Immunoblotting was performed using anti-GFP to detect AtDjB3–GFP, anti-Hsp101 as a marker for aggregated proteins, and anti-GAPDH as a loading control.

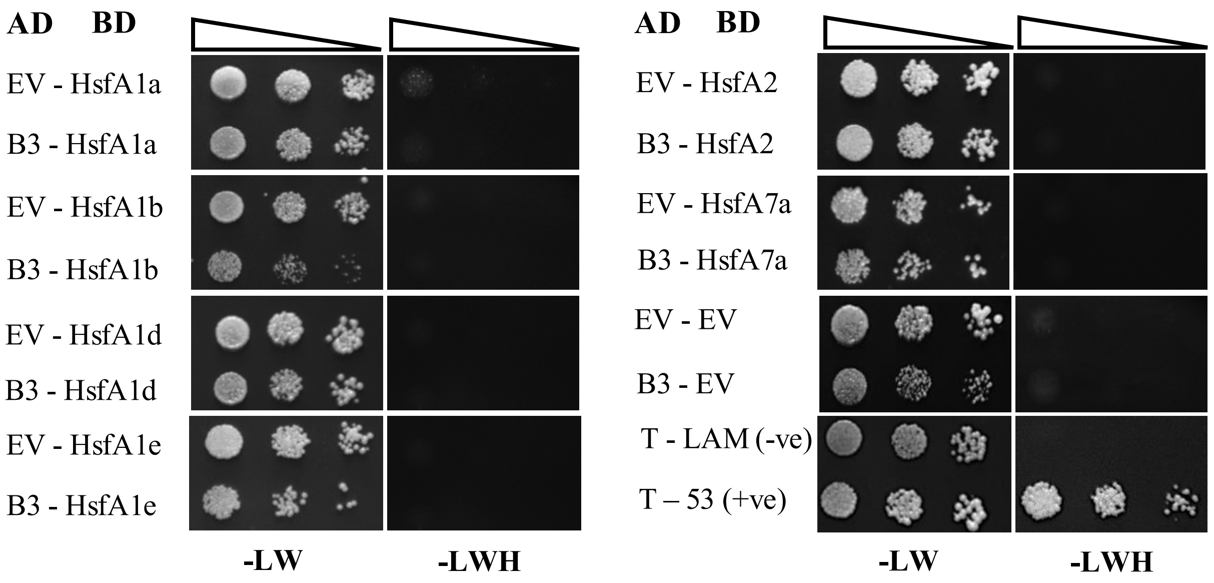


**Supplemental Figure 6. AtDjB3 does not interact with Arabidopsis Hsfs.** Yeast two-hybrid assay results showing co-expression of the indicated bait and prey combinations in Y2H gold strain yeast. Yeast was grown on non-selective (+His) or selective (–His) media. Growth on selective (–His) media indicates a protein-protein interaction. The positive control used was yeast expressing pGBKT7-53 (encoding Gal4 BD linked to murine p53) and pGADT7-T (encoding Gal4 AD fused with SV40 large T-antigen), which are known to interact. The negative control consisted of yeast expressing pGBKT7-Lam (Gal4 BD fused with Lamin) and pGADT7-T as well as yeast expressing empty vectors (BK-EV+AD-EV), both of which showed no interaction.

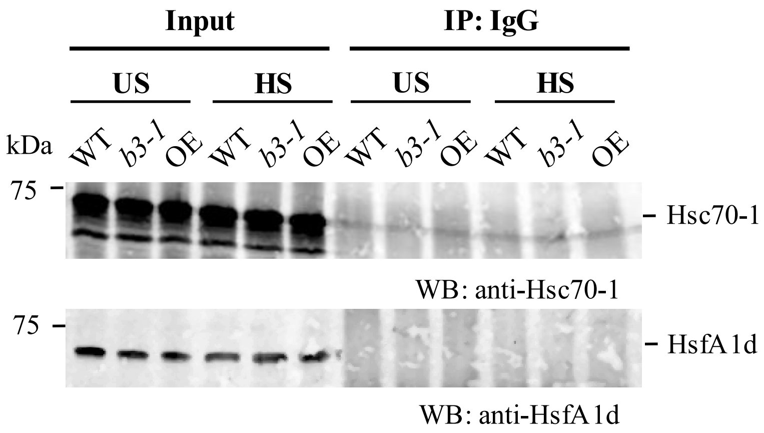


**Supplementary Figure 7. Negative control for co-immunoprecipitation.** Protein extracts from WT, *b3-1*, and *AtDJB3* overexpression seedlings under unstressed or heat-stressed conditions were subjected to co-immunoprecipitation using non-immune mouse IgG. Western blotting was performed with anti-Hsc70-1 and anti-HsfA1d antibodies.
